# PLANT NATRIURETIC PEPTIDE A antagonizes salicylic acid-primed cell death

**DOI:** 10.1101/592881

**Authors:** Keun Pyo Lee, Kaiwei Liu, Eun Yu Kim, Laura Medina-Puche, Jianli Duan, Yingrui Li, Haihong Dong, Ruiqing Lv, Zihao Li, Rosa Lozano-Duran, Chanhong Kim

## Abstract

Peptide hormones perceived in the cell surface via receptor proteins enable cell-to-cell communication and act in multiple biological processes through the activation of intracellular signaling. Even though Arabidopsis is predicted to have more than 1,000 secreted peptides, the biological relevance of the majority of these is yet to be established. Here, we demonstrate that PLANT NATRIURETIC PEPTIDE A (PNP-A), a functional analog to vertebrate atrial natriuretic peptides, antagonizes the salicylic acid (SA)-mediated cell death in the Arabidopsis *lesion-stimulating disease 1* (*lsd1*) mutant. While loss of PNP-A potentiates SA signaling, exogenous application of the PNP-A synthetic peptide or overexpression of PNP-A significantly compromises the SA-mediated cell death. Moreover, we identified a plasma membrane-localized receptor-like protein, which we name PNPAR (for PNP-A receptor), that binds PNP-A and is required to counteract SA responses. Our work identifies a novel peptide-receptor pair which modulates SA responses in Arabidopsis.

## Introduction

As plants are multicellular organisms, cell-to-cell communication is crucial for growth, development, and survival under ever-changing environmental conditions. This intercellular communication is largely mediated by secreted signals such as phytohormones, reactive oxygen species (ROS), small RNAs, and small peptide hormones^1^. While genome sequencing and transcriptome data analyses have identified more than 1,000 potential peptide hormones in *Arabidopsis thaliana*^2,3^, only a few of them are functionally characterized, and even fewer have a cognate receptor assigned^4–7^. Most peptide hormones undergo post-translational modifications (PTMs), such as proteolytic processing, glycosylation, formation of intra-disulfide bonds, proline hydroxylation, hydroxyproline arabinosylation, and sulfation of tyrosine residues, during or upon their secretion into the apoplastic space^3,5,7,8^. These modifications enable recognition by specific receptors, such as receptor-like kinases (RLKs)^9^ and receptor-like proteins (RLPs)^10^, on the surface of target cells, activating the relay of the signal into the cell interior^4–6,11^. In general, coupling of peptide-receptor results in transcriptional reprogramming, enabling the recipient cell to appropriately respond to an inbound factor following perception of the peptide secreted by the emitting cell. Like peptide hormones, the Arabidopsis genome also encodes hundreds of plasma membrane-associated RLKs and RLPs^9,10^. This diversity suggests a large number of potential peptide-receptor interactions that may facilitate the coordination and integration of multiple signaling pathways to modulate physiological processes following the perception of a broad range of external signals.

Plant natriuretic peptides (PNPs), functional analogs to vertebrate atrial natriuretic peptides (ANPs)^12^, are a novel type of peptide hormones that signal via guanosine 3’,5’-cyclic monophosphate (cGMP)^13–16^. In animals, the synthesis of cGMP from guanosine triphosphate (GTP) is catalyzed by natriuretic peptide receptors (NPRs), which possess protein kinase (PK) and guanylyl cyclase (GC) activities, following perception of ANPs^17^. Like ANPs, upon secretion to the apoplast, PNPs undergo formation of inter-disulfide bonds and proteolytic processing^15,17^. Although PNPs have been shown to affect a broad spectrum of physiological responses in plants, including stomata opening^14,18–20^, regulation of photosynthetic efficiency and photorespiration^19,21^, cellular water and ion (Ca^2+^, H^+^, K^+^ and Na^+^) homeostasis ^13,15,22^, increase in protoplast volume^14,15^, modulation of their own expression^23,24^ and resistance against biotic and abiotic stresses^20^, their mode of action is largely unclear and their cognate receptor proteins are unknown.

In the present study, we found that *PNP-A*, but not its close homologue *PNP-B*, is transcriptionally upregulated in the Arabidopsis *lesion-simulating disease 1* (*lsd1*) mutant prior to the onset of cell death, which requires NONEXPRESSER OF PR GENES 1 (NPR1), a key regulator of the salicylic acid (SA)-mediated signaling in plants^25,26^. The PNP-A processed (active) form is secreted into the apoplastic space and interacts with a previously uncharacterized plasma membrane-localized leucine-rich repeat RLP, named here PNPAR (for PNP-A Receptor), which is required for responses to PNP-A. While the lack of PNP-A or PNPAR potentiates the *lsd1*-conferred lesion-mimicking cell death, exogenous application or overexpression of PNP-A considerably compromises this response. In agreement with a role of PNP-A antagonizing SA responses, the exogenous treatment with this peptide results in increased susceptibility to a bacterial pathogen. Taken together, our results unveil a physiological function of the peptide hormone PNP-A as a negative modulator of SA signaling and identify PNPAR as its cognate receptor.

## Results

### The secreted PNP-A peptide antagonizes SA-dependent plant responses

We previously established a linear signaling pathway from SA to chloroplast-mediated programmed cell death, which largely contributes to the *lsd1*-conferred runaway cell death (hereafter *lsd1* RCD)^27^. The *lsd1* mutant, since its discovery in 1994^28^, has been utilized as a bio-tool to understand the molecular mechanisms underlying the regulation of cell death (especially the constraining mechanisms), because in this mutant cell death spontaneously increases in an uncontrolled manner^29–31^: several molecular components involved in SA-dependent signaling pathways, including NPR1, ENHANCED DISEASE SUSCEPTIBILITY 1 (EDS1), and PHYTOALEXIN DEFICIENT 4 (PAD4), have been identified as required for the *lsd1* RCD^27,32,33^. Recently, we carried out a global transcriptome analysis that revealed a substantial number of genes rapidly upregulated before the onset of the *lsd1* RCD^27^. Among them, we identified the transcript of the PNP-A peptide hormone, of which the mode of action and the cognate receptor remain to be elucidated in plants. Unlike the upregulation of *PNP-A* (Supplementary Fig. 1a,b), the transcript of *PNP-B*, a *PNP-A* homolog, was undetectable in *lsd1* as well as wild-type (WT) plants, indicating the specificity of PNP-A toward the *lsd1* RCD. Because SA signaling primes the *lsd1* RCD^27^ and *PNP-A* belongs to a group of SA-responsive genes^34^, next we examined whether SA and its key signaling component NPR1 regulate *PNP-A* expression. While *PNP-A* was clearly induced in WT plants upon SA treatment, it was not in *npr1* (Supplementary Fig. 1c), indicating that SA and its bona fide receptor NPR1 act to positively regulate the expression of *PNP-A*. Alongside with the notable attenuation of the *lsd1* RCD^27^, the loss of NPR1 in the *lsd1* background completely abrogated *PNP-A* expression (Supplementary Fig. 1d).

To explore the potential causal relationship between the rapid upregulation of *PNP-A* and the development of *lsd1* RCD, the plant phenotype resulting either from inactivation or overexpression of PNP-A was examined. For this, a *pnp-A* knockout mutant was crossed with *lsd1* to create the *lsd1 pnp-A* double mutant plants, and two independent *lsd1* transgenic plants overexpressing *PNP-A* under the control of the *CaMV 35S* (35S) promoter were chosen based on the increased levels of *PNP-A* transcripts as compared to *lsd1* (Supplementary Fig. 2a,b). The genetic inactivation of PNP-A significantly facilitated the emergence of RCD, while its overexpression drastically attenuated the phenotype (Fig. 1a; Supplementary Fig. 3). The degrees of foliar cell death and chlorosis were evaluated by examining ion leakage and maximum photochemical efficiency of photosystem II (Fv/Fm), respectively (Fig. 1b,c). Given that overexpression of PNP-A significantly compromised the *lsd1* RCD, which is largely dependent on SA signaling, we anticipated that PNP-A might antagonize the SA-primed intercellular signaling. Reverse transcription quantitative PCR (RT-qPCR) to examine the expression level of genes involved in SA synthesis, signaling and/or responses, including *ISOCHORISMATE SYNTHASE 1* (*ICS1*), *EDS1, PAD4*, *PATHOGENESIS RELATED* (*PR*)*1*, and *PR2*, revealed that PNP-A overexpression markedly repressed the *lsd1*-induced SA signaling (Supplementary Fig. 4). Based on this, we hypothesized that, upon secretion to the apoplastic space, PNP-A diffuses to adjacent cells and interacts with its cognate receptor to activate intracellular signaling, ultimately antagonizing SA-primed cell death responses. Even though PNP-A was previously suggested to be a secreted peptide hormone^24^, its presence in the apoplastic space has not been previously detected. To test this idea, we decided to examine the subcellular localization of PNP-A fused to the GREEN FLUORESCENT PROTEIN (GFP) upon its transient expression driven by the 35S promoter in *Nicotiana benthamiana* leaves. Besides its post-translational modification (i.e. intra-disulfide bond formation between Cys42 and Cys65) in the apoplast, it was proposed that the PNP-A precursor is proteolytically processed to its active form, resembling what happens to the vertebrate ANP^35,36^. To avoid the cleavage of GFP, we fused the GFP with the truncated PNP-A (tPNP-A, Met1 to Tyr69) containing an N-terminal signal peptide (SP, Met1 to Lys29) and the putative active domain (Pro36 to Tyr69). The obtained confocal images clearly demonstrate that the tPNP-A-GFP protein is exclusively localized in the apoplastic space (Fig. 1d).

**Figure 1.**
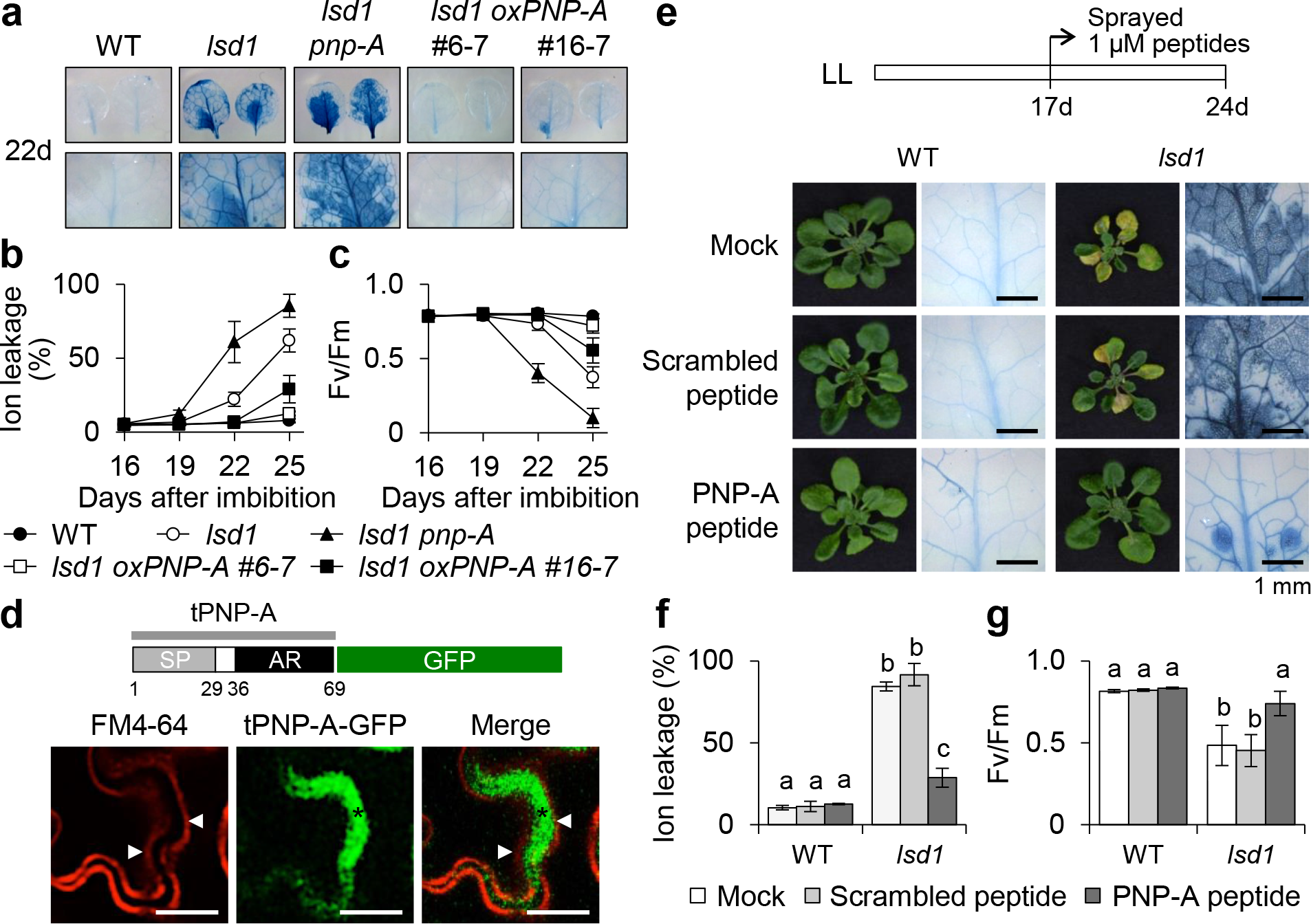
PNP-A acts to repress *lsd1* RCD. (**a**) Twenty two-days (d)-old plants of wild-type (WT), *lsd1*, *lsd1 pnp-A*, and two independent 35S:*PNP-A lsd1* transgenic lines (#6-7, #16-7) grown under continuous light condition (CL) were collected to examine leaf RCD. First row: the RCD phenotype in the first or second leaves from the each genotype was visualized by trypan blue (TB) staining. Second row: the images of the first row were enlarged. (**b**,**c**) For measurements of ion leakage (**b**) and maximum photochemical efficiency of PSII (Fv/Fm) (**c**), first or second leaves from plants were harvested at the indicated time points. Ten leaves per genotype were used for each measurement. Data represent the means from three independent measurements. Error bars indicate standard deviation. (**d**) PNP-A is localized to the apoplastic space. Localization of the truncated PNP-A (tPNP-A, Met1 to Tyr69), including signal peptide (SP) and active region (AR), fused with GFP (PNPA-GFP) upon transient expression in *N. benthamiana* leaves. FM4–64 was used to stain plasma membrane (PM). Cell plasmolysis was performed by treatment of 0.8 M mannitol for 30 min. An asterisk indicates the apoplastic space formed by the shrinking protoplast, and triangles indicate the retracted PM. Scale bar = 20 μm. (**e-f**) Seventeen-day-old WT and *lsd1* plants grown under CL were treated with water (Mock) or 1 μM scrambled or active form of AtPNP-A synthetic peptide. After 7 days of the treatment, the relative levels of foliar RCD were determined by TB staining (**e**), ion leakage (**f**) and Fv/Fm measurements (**g**). Data represent the means from three independent measurements. Error bars indicate standard deviation.

The antagonizing impact of PNP-A on the *lsd1* RCD was further examined via a pharmacological approach by using synthesized active and dormant (scrambled) PNP-A peptides (for details on the synthetic peptides, see Supplementary Fig. 5). While no impact of the scrambled peptide was observed, the *lsd1* mutant plants treated with the active form of the PNP-A peptide exhibited drastically reduced RCD (Fig. 1d-f). Next, the impact of SA on plants deficient for or overexpressing PNP-A was analyzed. Exogenous application of high dosage of SA is known to greatly inhibit plant growth because of a trade-off effect^37,38^, i.e. an enhanced immune response limits plant growth or *vice versa*. It was clear that the negative impact of SA on plant growth was remarkably potentiated in *pnp-A* mutant plants, whereas it was less obvious in PNP-A overexpressing plants relative to WT (Fig. 2a,b), again pointing at a function of PNP-A in restricting SA responses.

**Figure 2.**
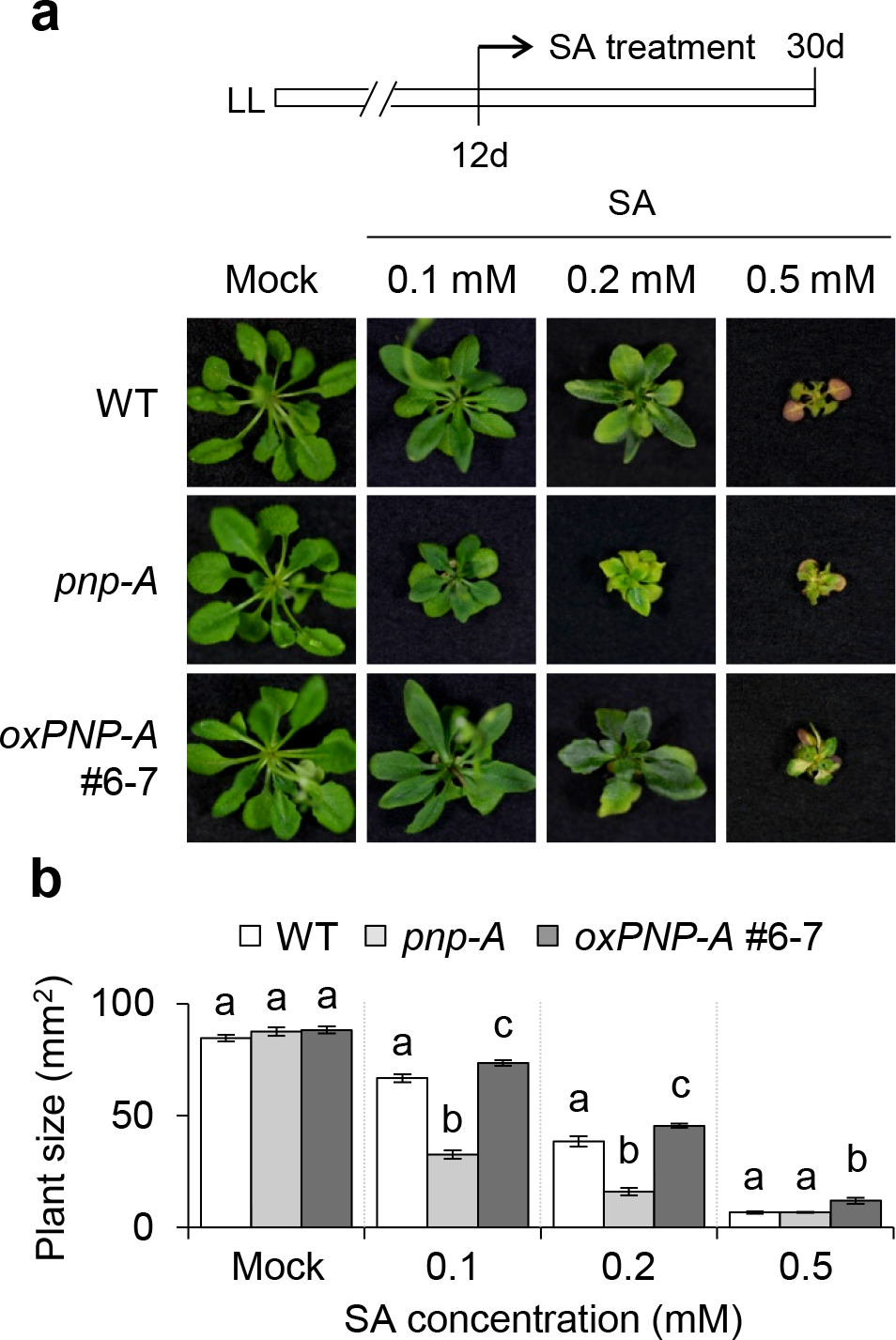
PNP-A is required to counteract SA-mediated growth inhibition. Twelve-day-old plants of WT, *pnp-A* and *p35S:PNP-A lsd1* (#6-7) grown on MS medium under CL were transferred to fresh MS medium containing the different concentrations of SA as indicated. The representative foliar phenotype (**a**) and plant size (**b**) of each genotype were shown. For the measurement of plant size, fifteen plants per genotype were used for each measurement. Data represent the means from three independent measurements. Error bars indicate SD. Lowercase letters indicate statistically significant differences between mean values (*P* < 0.05, one-way ANOVA with post-hoc Tukey’s HSD test).

### The PNP-A peptide physically interacts with the PM-localized LRR family protein PNPAR

The vertebrate PNP analogues interact with GC-coupled protein receptors which catalyze the conversion of GTP into cGMP upon binding of the peptide^17^. The resulting increased cellular cGMP acts as a second messenger in the intracellular signal transduction mostly by activating kinases, which is implicated in ion channel conductance, glycogenolysis, relaxation of smooth muscle tissues, and inhibition of platelet aggregation^39^. In plants, cGMP is also a crucial signaling molecule involved in stress responses^16,40,41^, ion homeostasis through the regulation of cyclic nucleotide gated channels^42,43^, nitric oxide (NO)-dependent signaling^44,45^, hormonal signaling^46,47^, and phytochrome-dependent transcriptional regulation^48^. Importantly, it was previously reported that PNP-A interacts with a novel leucine-rich repeat (LRR) protein, namely PNP-R1, which is predicted to contain a putative N-terminal SP, a LRR N-terminal (LRRNT) domain, a transmembrane (TM) domain, two LRR domains, and a PK domain followed by a GC catalytic center at the C-terminus^15^. Since the TM domain is located between the LRRNT and the LRR domains, this prediction implies that the LRR, PK and GC domains face the cytosol, whereas the LRRNT domain protrudes toward the extracellular space, or vice versa. However, to our surprise, when the deduced amino acid sequence of the PNP-R1 was subjected to a search for conserved domains by using several bioinformatics tools, including TMHMM server v. 2.0^49^, Phobius program^50^, NCBI’s Conserved Domain Database^51^ and InterPro^52^, the TM domain was not identified. Moreover, neither the PK nor the GC catalytic domains were predicted. Given that several plant GCs have low sequence homology with the annotated GCs of other organisms^53^, the GC catalytic center of PNP-R1 had been identified on the basis of non-canonical sequences^15^. While it was shown that PNP-R1 possesses *in vitro* GC activity^15^, so far there is no experimental evidence for its kinase activity.

In our system (i.e. *lsd1* mutant background), we could not identify the GC-containing receptor protein as a putative PNP-A-interacting protein through pull-down with biotin-conjugated PNP-A peptide coupled to mass spectrometry analysis (Supplementary Table 1). Instead, we found a typical receptor-like protein, which is predicted to contain a SP, an LRRNT domain followed by nine LRRs, and a TM domain, but lacking cytosolic activation domain (e.g. kinase) (Fig. 3a). Here, we tentatively named this RLP PNP-A receptor protein (PNPAR, At5g12940). The fluorescent signal (red) of the FM4-64 dye co-localizes with the florescence signal (yellow) of PNPAR-YFP, and both signals were retained in the plasma membrane (PM) after plasmolysis (Fig. 3b), indicating that PNPAR is localized to PM. Biomolecular fluorescence complementation (BiFC), *in vitro* pull-down, and co-immunoprecipitation analysis in *N. benthamiana* leaves substantiated the direct interaction between PNP-A and PNPAR (Fig. 3c-e).

**Figure 3.**
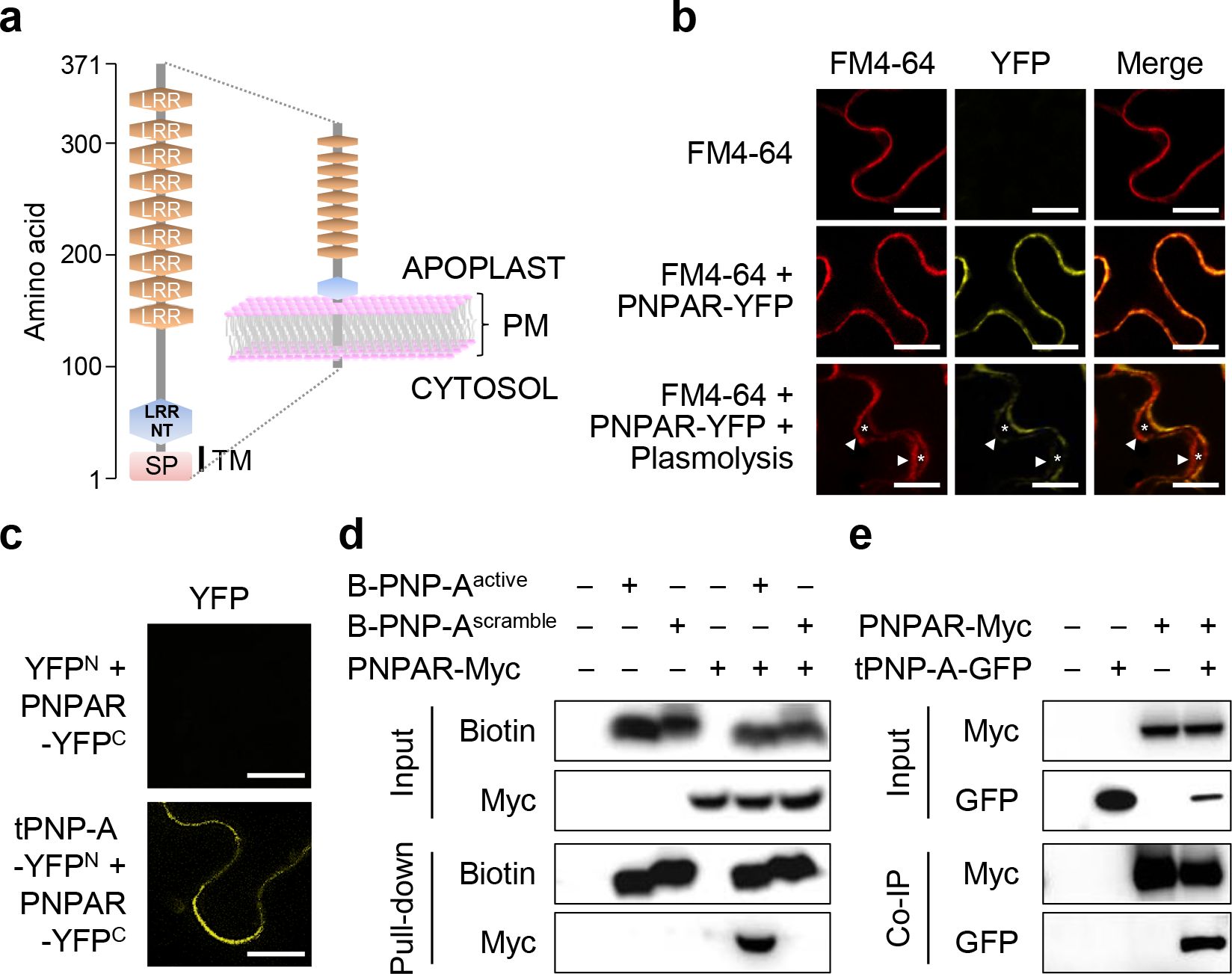
PNP-A interacts with a novel plasma membrane-localized LRR receptor-like protein. (**a**) Schematic illustration of the predicted domain structure and topology of the putative PNP-A-interacting LRR receptor, namely PNPAR. The PNPAR contains the extracellular domains consisting of 9 LRRs and a LRRNT. SP indicates a signal peptide present at the N-terminus of PNPAR and is responsible for targeting to plasma membrane (PM). TM marks the predicted transmembrane domain. (**b**) YFP fluorescence was observed in the PM when the YFP-tagged PNPAR (PNPAR-YFP) was expressed in *N. benthamiana* leaves. FM4–64 was used to stain PM. Cell plasmolysis was performed by treatment of 0.8 M mannitol for 30 min. In plasmolyzed cells, asterisks indicate the apoplastic space formed by the shrinking protoplast, and triangles indicate the retracted PM. Scale bar = 20 μm. (**c**) Interaction of PNP-A with PNPAR by biomolecular fluorescence complementation (BiFC) assay. YFP fluorescence was observed in the PM when the N-terminal part of YFP tagged with the truncated PNP-A (tPNP-A, Met1 to Tyr69 including SP and active region) (tPNP-A-YFP^N^) was coexpressed with the C-terminal part of the YFP tagged with PNPAR (PNPAR-YFP^C^) in *N. benthamiana* leaves. Scale bar = 20 μm. (**d**) *In vitro* pull-down assay of N-terminally biotinylated active region of PNP-A synthetic peptide (B-PNP-A^active^) with PNPAR fused with Myc tag (PNPAR-Myc) upon transient expression in *N. benthamiana* leaves. N-terminally biotinylated-scrambled PNP-A peptide (B-PNP-A^scrambled^) was used as a negative control as it does not interact with PNPAR. (**e**) Co-immunoprecipitation (Co-IP) of tPNP-A-GFP with PNPAR-Myc upon transient coexpression in *N. benthamiana* leaves.

### PNPAR is required for the PNP-A-mediated repression of SA responses

To further corroborate that PNPAR is a bona fide receptor of PNP-A, two knockout mutant alleles of *PNPAR* (*pnpar-1* and *pnpar-2*; Fig. 4a) were crossed with *lsd1* plants to create two independent *pnpar lsd1* double mutants. The genetic inactivation of *pnpar* in *lsd1* revealed a probable genetic interaction between PNP-A and PNPAR, which was evident from the potentiated foliar cell death and chlorosis as well as the significant decrease of Fv/Fm in the both *lsd1 pnpar-1* and *lsd1 pnpar-2* double mutant plants as compared to *lsd1* (Fig. 4b-e). The enhanced RCD phenotype in *pnpar lsd1* plants was accompanied by the heightened expression of *PR1* and *PR2* (Fig. 4f), implying that intracellular immune responses seem to be reinforced in the absence of PNPAR. By contrast, a *lsd1 pnp-r1* double mutant showed equivalent degrees of RCD and expression of *PR1* and *PR2* as compared to those of *lsd1* (Fig. 4b-f), suggesting that PNP-R1 is not involved in the *lsd1* RCD or immune responses. In addition, unlike *lsd1* and *lsd1 pnp-A*, in which the active PNP-A synthetic peptide led to the attenuation of the *lsd1* RCD, the *lsd1 pnpar* double mutant plants were insensitive to this treatment (Fig. 5a-c). Conversely, *pnpar* mutant plants exhibited an extreme sensitivity to exogenously applied SA, like *pnp-A* (Fig. 5d,e), as shown by the drastic growth inhibition and chlorosis, supporting the biological function of the PNP-A/PNPAR pair in curtailing SA responses.

**Figure 4.**
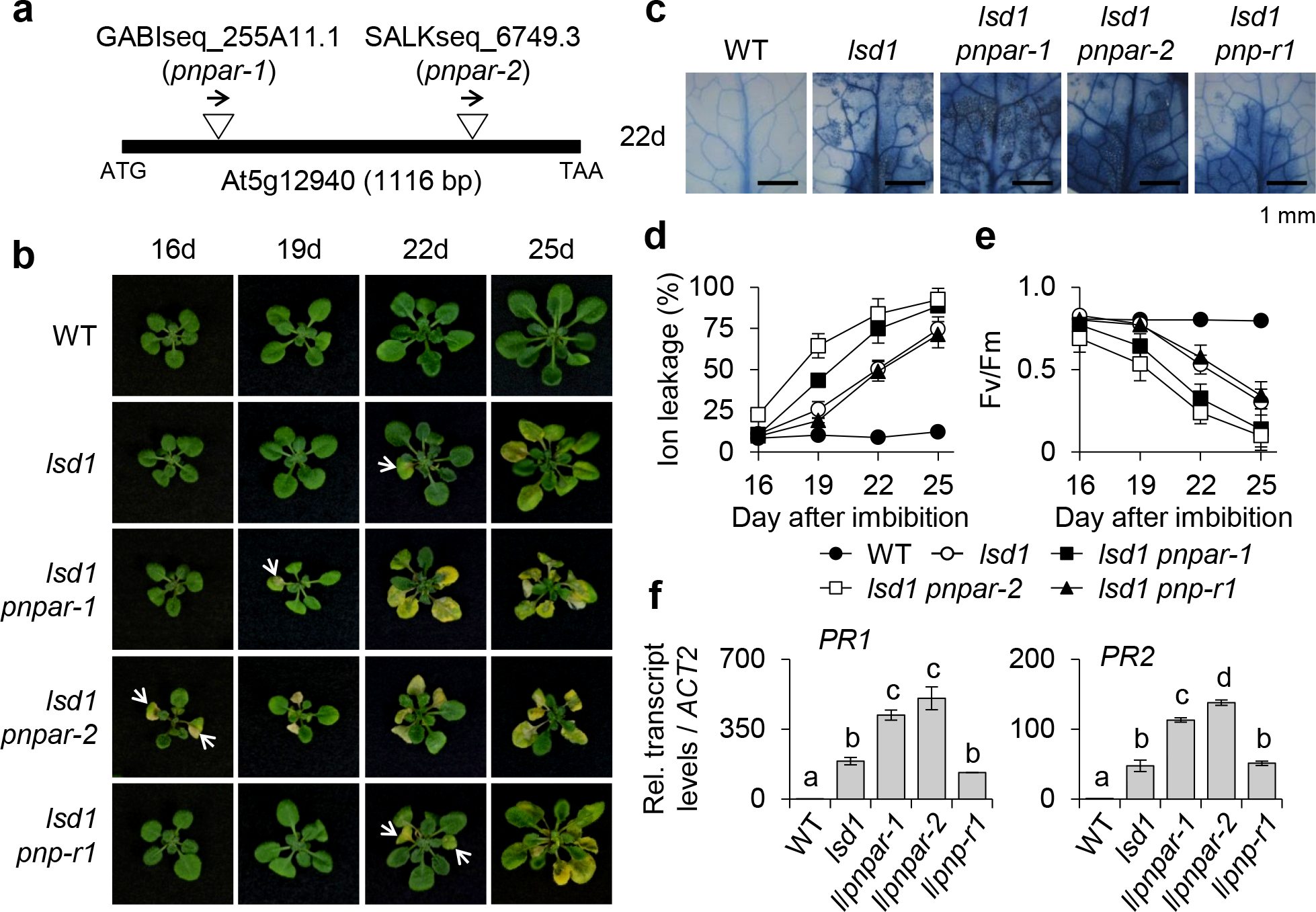
PNPAR is required to counteract SA-mediated *lsd1* RCD and immune responses. (**a**) The insertion positions of T-DNAs in two *Arabidopsis pnpar* mutant alleles, *pnpar-1* and *pnpar-2*. It should be noted that PNPAR does not contain any intron. (**b**) WT, *lsd1*, *lsd1 pnpar-1*, *lsd1 pnpar-2* and *lsd1 pnp-r1* plants were grown under CL and the emergence and the spread of RCD were monitored at the indicated time points. The images of representative plants are shown at the same scale. (**c**) The degree of RCD in the leaves from each genotype grown under CL for 22 days was visualized by TB staining. (**d,e**) For measurements of ion leakage (**d**) and Fv/Fm (**e**), first or second leaves from each genotype grown under CL were harvested at the indicated time points. Ten leaves per genotype were used for each measurement. Data represent the means from three independent measurements. Error bars indicate standard deviation (SD). (**f**) The relative expression levels of *PR1* and *PR2* were determined using qRT-PCR. *ACT2* was used as an internal standard. The data represent the means of three independent biological replicates. Error bars indicate SD. Lowercase letters indicate statistically significant differences between mean values (*P* < 0.01, one-way ANOVA with post-hoc Tukey’s HSD test).

Recently, the response of plants overexpressing *Arabidopsis* PNP-A to a bacterial pathogen was examined: the results show that PNP-A potentiates the expression of defense-related genes, including *PR1*, conferring increased resistance to *Pseudomonas syringae* pv. *tomato* (*Pst*) DC3000 infection^20^. This is, however, inconsistent with our finding that PNP-A negatively regulates the expression of *PR* genes along with other SA-responsive genes in *lsd1*. For such reason, we decided to examine whether PNP-A affects host resistance against *Pst* DC3000 in our experimental system. As shown in Fig. 6a, exogenous application of the PNP-A synthetic peptide, but not of the scrambled peptide, enhanced susceptibility to *Pst* DC3000, providing another evidence that PNP-A negatively regulates plant defense responses. Moreover, the *pnpar* mutant was insensitive to the PNP-A synthetic peptide, unlike the *pnp-r1* mutant (Fig. 6b). Taken together, our results strongly suggest that the PNP-A/PNPAR-mediated intracellular signaling counteracts SA responses, which has a pivotal role in the modulation of defense responses^54^.

**Figure 5.**
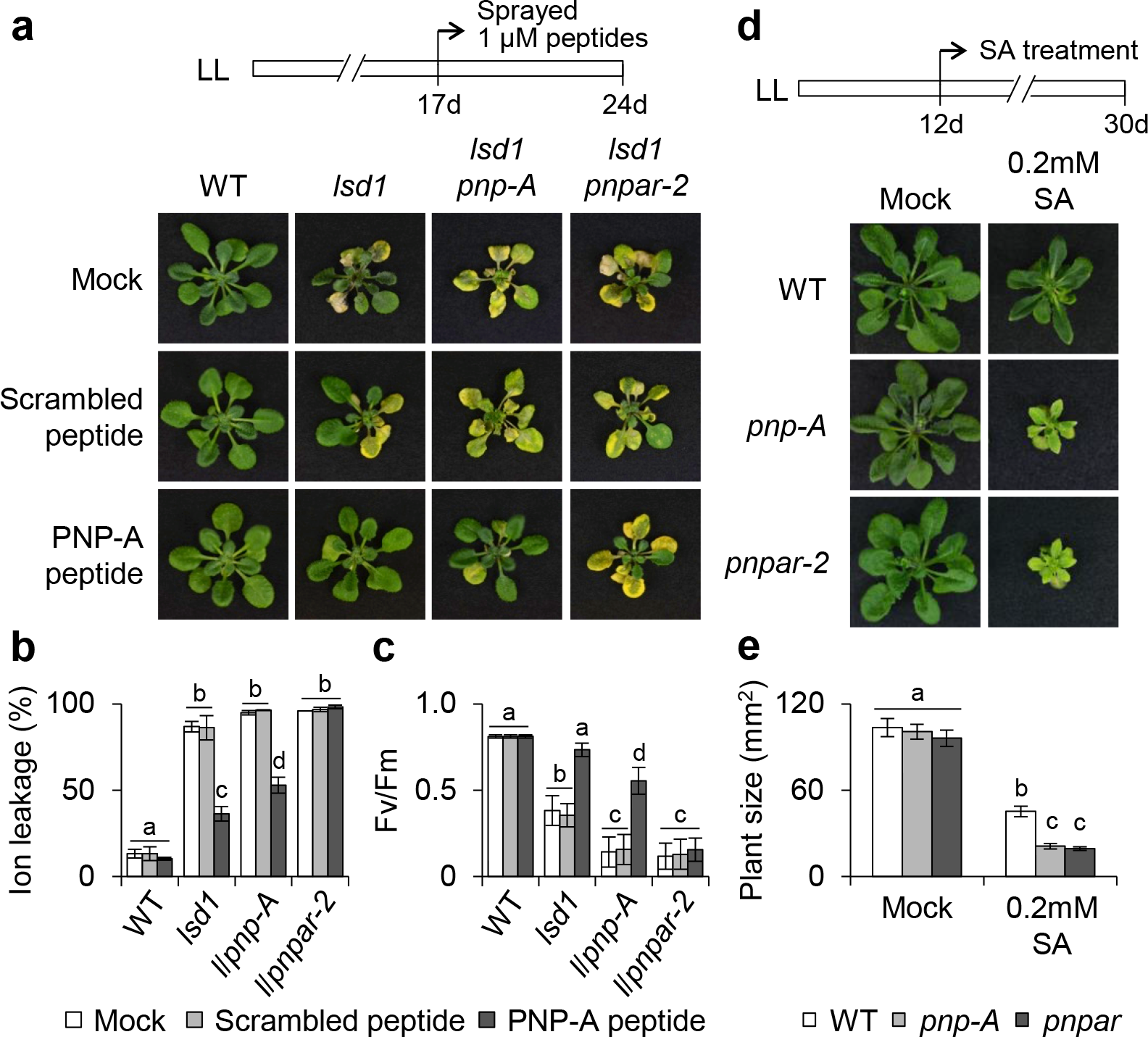
*pnpar* mutant plants are insensitive to PNP-A and hypersensitive to SA. (**a-c**) Seventeen-day-old plants of WT, *lsd1*, *lsd1 pnp-A* and *lsd1 pnpar-2* grown under CL were treated with PNP-A or scrambled peptides and kept for 7 days under CL. Afterward, the RCD phenotype (**a**), ion leakage (**b**) and Fv/Fm (**c**) were examined. The representative images are shown at the same scale. For the measurements of ion leakage and Fv/Fm, ten leaves per genotype were used for each measurement. Data represent the means from three independent measurements. Error bars indicate standard deviation (SD). (**d**,**e**) Twelve-day-old plants of WT, *pnp-A* and *pnpar-2* grown on MS medium under CL were transferred to MS medium in the absence (Mock) or presence of 0.2mM SA and kept for 18 days under same growth condition. The representative foliar phenotype (**d**) and plant size (**e**) of each genotype were shown. For the measurement of plant size, fifteen plants per genotype were used for each measurement. Data represent the means from three independent measurements. Error bars indicate SD. Lowercase letters in **b**, **c** and **e** indicate statistically significant differences between mean values (*P* < 0.05, one-way ANOVA with post-hoc Tukey’s HSD test).

## Discussion

The *PNP-A* gene is transcriptionally upregulated in response to abiotic stresses including UV-B, salt, osmotic, nutrient deficiencies, and ozone^55^, indicating that PNP-A may modulate plant responses to a multitude of environmental factors. We found that *PNP-A* was also upregulated in *lsd1* mutant plants prior to the onset of RCD (Supplementary Fig. 1a,b) which is known to be spread in an uncontrolled manner by various biotic and abiotic factors, such as excess light, red light, UV radiation, root hypoxia, cold and bacterial infection^28,32,56–60^.

Since this *lsd1* RCD is mediated by molecular components, such as EDS1, PAD4 and NPR1, involved in SA accumulation and the SA-dependent systemic acquired resistance (SAR) pathway^27,32,33^, it is reasonable to assume that PNP-A-mediated intercellular signaling participates in the regulation of the spread of cell death in *lsd1* that results from inappropriately induced SA-dependent plant defense/immune responses. In fact, a large-scale co-expression analysis indicates that PNP-A is highly co-expressed with genes associated with the SAR pathway^55^. A previous proteomic analysis of plant cells treated with synthetic Arabidopsis PNP-A peptide also demonstrates that PNP-A affects the abundance of proteins involved in cellular oxidation-reduction processes and in responses to biotic and abiotic stresses^61^.

In this study, we reveal a molecular pathway by which the PNP-A peptide hormone negatively regulates SA-mediated plant immune responses. Upon SA- and NPR1-dependent transcriptional upregulation of *PNP-A* (Supplementary Fig. 1c,d), the apoplastic PNP-A peptide physically interacts with its PM-localized cognate receptor protein PNPAR (Fig. 3). The PNP-A/PNPAR pair acts to inhibit SA signaling, antagonizing the SA-triggered RCD in the *lsd1* mutant (Figs 1 and 4), as well as the SA-dependent growth retardation (Figs 2 and 5) and increasing the plant susceptibility to a virulent bacterial pathogen (Fig. 6). Another kind of intercellular signaling molecules, ROS produced by PM-associated Arabidopsis respiratory bust oxidase (AtRBOH) family proteins, can also antagonize SA-dependent signaling to restrict the spread of cell death in the *lsd1* mutant and upon infection of avirulent bacterial pathogens^62^. Therefore, both of these systems may play an important role in fine-tuning plant immune responses to avoid inappropriate induction of SA-dependent death signals in cells spatially separated from infected or damaged cells, thereby minimizing tissue damage.

It has to be noted that the RLP PNPAR lacks an intracellular domain with enzymatic activity (Fig. 3a). Because RLPs frequently act coordinately with other LRR proteins harboring intracellular signaling domains to perceive extracellular peptide signals and instigate intracellular signaling ^11,63,64^, we hypothesize that PNPAR may form a complex with a co-receptor protein, e.g. an LRR-RLK, to relay the signal upon recognition of the PNP-A peptide. Therefore, finding the potential co-receptor of PNPAR will be essential to unveil downstream signaling components, paving the way for the eventual full dissection of the PNP-A signaling pathway.

**Figure 6.**
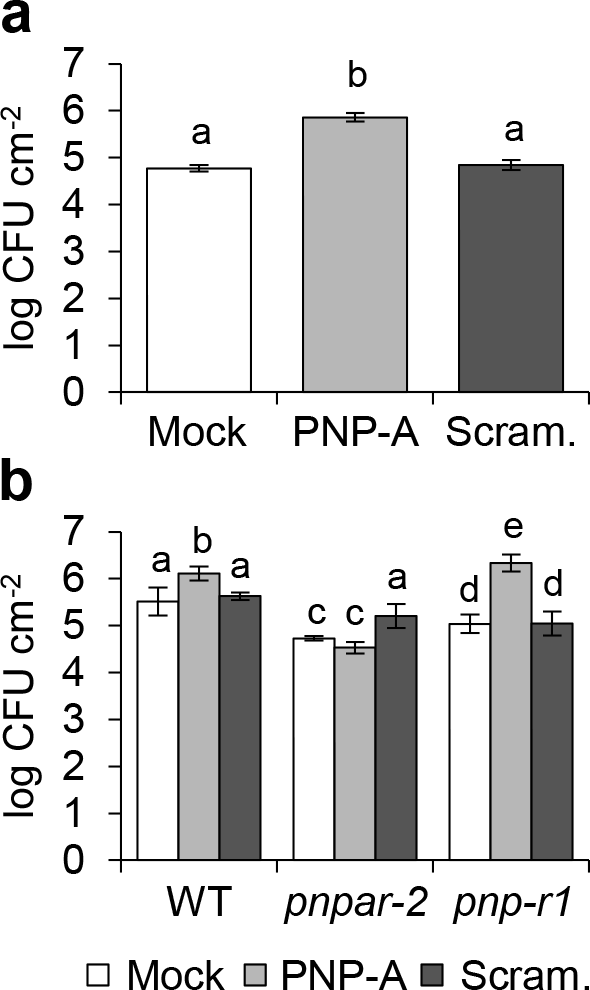
The PNP-A peptide enhances plant susceptibility to *P. syringae* pv. *tomato* DC3000 in a PNPAR-dependent manner. (**a,b**) Five-week-old short-day-grown plants of WT (**a**) and WT, *pnpar-2* and *pnp-r1* (**b**) were pre-treated with 5 μM PNP-A synthetic peptide (PNP-A) or scrambled peptide (Scram.) control 6 h prior to bacterial infiltration. Three days later, bacteria were extracted from three different leaves of 4 independent plants and incubated at 28 °C for 2 days to evaluate growth. Bars represent SE of n = 4. Lowercase letters indicate statistically significant differences between mean values (*P* < 0.001, one-way ANOVA with post-hoc Bonferroni’s multiple comparison test). This experiment was repeated twice with similar results. CFU: colony-forming unit.

## Methods

### Plant materials and growth conditions

All the *Arabidopsis* genotypes used in this study are Columbia (Col-0) ecotype. *Arabidopsis* mutant seeds of *lsd1-2* (SALK_042687)^27^, *pnp-A* (SALK_000951)^15,20^, *npr1* (SALK_204100), *pnpar-1* (GABIseq_255A11.1), *pnpar-2* (SALKseq_6749.3), and *pnp-r1* (GABI-KAT_180G04)^15^ were obtained from the Nottingham Arabidopsis Stock Centre (NASC). We generated and genotyped double mutants by crossing homozygous single mutant plants and using appropriate primers (Supplementary Table 2). Seeds were surface sterilized by soaking in 1.6 % hypochlorite solution for 10 min, followed by washing five times with sterile water. Seeds were then plated on Murashige and Skoog (MS) medium (Duchefa Biochemie) containing 0.65% (w/v) agar (Duchefa Biochemie). After a three-day stratification at 4°C in darkness, seeds were placed in a growth chamber (CU-41L4; Percival Scientific) under continuous light condition. The light intensity was maintained at 100 μmol·m^−2^·s^−1^ at 22 °C ± 2°C. For pathogen infection assays, plants were grown on jiffy pellets in a controlled-environment chamber under short-day conditions (8 h light/16 h dark cycle) at 20-22 °C.

### Generation of *PNP-A* overexpression lines

The stop codon-less *PNP-A* coding sequence (CDS) was cloned into a pDONR221 Gateway vector (Thermo Scientific) via a Gateway BP reaction (Thermo Scientific) and subsequently recombined into the Gateway-compatible plant binary vector pGWB651^65^ via a Gateway LR reaction (Thermo Scientific). The generated vector was transformed into the *Agrobacterium tumefaciens* strain GV3101 using the heat shock method. After generating *Arabidopsis* transgenic plants in WT background using *Agrobacterium*-mediated transformation by the floral dip method^66^, homozygous T3 transgenic plants were selected on MS medium containing 12.5mg/l Basta (Sigma).

### RNA extraction and reverse transcription quantitative PCR (RT-qPCR)

Total RNA (1ug) extracted from foliar tissues, using the Spectrum Plant Total RNA Kit (Sigma-Aldrich) was reverse-transcribed with the HiScript II Q RT SuperMix for qPCR (Vazyme Biotech) according to the manufacturer’s recommendations. The RT-qPCR was conducted in triplicates on a QuantStudio™ 6 Flex Real-Time PCR System (Applied Biosystems) with SYBR Green Master Mix (Vazyme Biotech). Relative transcript levels were calculated by the ddCt method^67^ and normalized to the *ACTIN2* (At3g18780) transcript level. The sequences of the primers used in this study are listed in Supplementary Table 2.

### Bimolecular fluorescence complementation (BiFC) assay

BiFC assays were performed with a split-YFP system in *N. benthamiana* leaves as described previously^68^. In brief, the pDONR/Zeo entry vectors (Thermo Scientific) containing the 207 bp fragment of *PNP-A* CDS (counted from start codon) and the stop codon-less full CDS of *PNPAR* were recombined into the split-YFP vectors pGTQL1211YN and 1221YC, respectively, through a Gateway LR reaction (Thermo Scientific). For the BiFC assay, *A. tumefaciens* mixtures carrying the appropriate BiFC constructs were infiltrated with a 1 mL needle-less syringe into the abaxial side of 4-week-old *N*. *benthamiana* leaves. After 72 h, the presence of YFP signal was evaluated with a Leica TCS SP8 SMD (Leica Microsystems).

### Co-immunoprecipitation (Co-IP) assay

For Co-IP assay, the pDONR/Zeo entry vector containing the 207 bp fragment of *PNP-A* CDS (*tPNP-A*; counted from start codon) or the stop codon-less full CDS of *PNPAR* was recombined into the destination vector pGWB651 for C-terminal fusion with GFP or pGWB617 for C-terminal fusion with 4xMYC through Gateway LR reaction (Thermo Scientific) to create *p35S:tPNP-A-GFP* and *p35S:PNPAR-4xMYC*. One or both of the vectors were expressed alone or co-expressed in 4-week-old *N. benthamiana* leaves after *Agrobacterium* infiltration. Total protein was extracted with IP buffer [50 mM Tris-HCl (pH 7.5), 150 mM NaCl, 10% glycerol,1.0 mM EDTA, 1% Triton X-100, 1mM Na_2_MoO_4_·2H_2_O, 1mM NaF, 1.5mM Na_3_VO_4_, 1mM PMSF and 1x cOmplete protease inhibitor cocktail (Roche)]. After protein extraction, 20 μL of Myc-Trap magnetic agarose beads (Myc-TrapMA, Chromotek) were incubated with 40 mg of the total protein extract for 12 h at 4 °C by vertical rotation. The beads were washed five times with washing buffer (IP buffer with 0.1% Triton X-100) and then eluted with 2x SDS protein sample buffer for 10 min at 95 °C. The eluates were subjected to 10% (w/v) SDS-PAGE gels and the interaction between co-expressed proteins determined by immunoblot analysis using a mouse anti-MYC monoclonal antibody (1:10,000; Cell Signaling Technology) and a mouse anti-GFP monoclonal antibody (1:5,000; Roche).

### Preparation of peptides

N-terminal biotin-labeled PNP-A peptide and its scrambled peptide control were synthesized and purified to > 95 % purity on a high-pressure liquid chromatography (HPLC) by Sangon biotech (China). Peptides were dissolved in 20% acetic acid (1.5 mM stock solution) and diluted with distilled water to the desired concentrations before use. The amino acid sequences of peptides are shown in Supplementary Fig. 5.

### Peptide pull-down assay and mass spectrometry analysis

Twenty-day-old *lsd1* mutant plants grown under CL on MS medium were harvested and homogenized to a fine power in liquid nitrogen using a mortar and pestle. Approximately 10 g of fine power were used to extract total protein with the IP buffer as mentioned above. After protein extraction, 100ul of Dynabeads M-280 Streptavidin beads (Thermo Scientific) prebound with or without the N-terminal biotinylated PNP-A peptide or its scrambled peptide, were incubated with 100 mg of the total protein extract for 12 h at 4 °C by vertical rotation. The beads were washed five times with the washing buffer (IP buffer with 0.1% Triton X-100) and then washed two times with 1x PBS buffer. After protein elution from the beads with SDT-lysis buffer [100mM Tris-HCl (pH7.6), 4% (w/v) SDS, 0.1M DTT], the eluates were subjected to mass spectrometry analysis as described previously^69^.

Pull-down assays were also conducted with *N. benthamiana* leaves harboring the *p35S:PNPAR-4xMYC* after *Agrobacterium* infiltration. Total protein was extracted with the IP buffer from 5 g of *N. benthamiana* leaves. After pull-down assay as described above, the eluates were subjected to NuPAGE Bis-Tris 4-12% protein gel (Thermo Scientific), and the interaction between PNPAR and the PNP-A synthetic peptide was determined by immunoblot analysis using a mouse anti-MYC monoclonal antibody (1:10,000; Cell Signaling Technology) and a mouse anti-biotin monoclonal antibody (1:2,000; Sigma).

### Subcellular localization and confocal laser-scanning microscopy (CLSM)

The pDONR/Zeo entry vectors (Thermo Scientific) containing stop codon-less CDS of *PNPAR* was recombined into the destination vector pGWB641 for C-terminal fusion with YFP through a Gateway LR reaction (Thermo Scientific) to create *p35S:PNPAR-YFP*. To determine the subcellular localization of PNPAR and tPNP-A, the *p35S:PNPAR-YFP* and *p35S:tPNP-A-GFP* constructs were transformed into *A. tumefaciens* strain GV3101 and transiently expressed in *N. benthamiana* leaves. The GFP, YFP, and FM4-64 fluorescence signals were detected by CLSM analysis using Leica TCS SP8 SMD (Leica Microsystems) 72 hours after infiltration. All the images were acquired and processed using the Leica LAS AF Lite software version 2.6.3 (Leica Microsystems). Cell plasmolysis was performed by treatment with 0.8 M mannitol for 30 min.

### Determination of cell death

Cell death was visualized and quantified using trypan blue (TB) staining and electrolyte leakage measurements as previously described^27^. Briefly, first or second leaves from plants grown under CL on MS medium were submerged in TB staining solution (10 g phenol, 10 ml glycerol, 10 ml lactic acid and 0.02 g trypan blue in 10 ml H_2_O), diluted with ethanol 1:2 (v/v), and boiled for two min on a water bath. After a 16 h-incubation at room temperature on a vertical shaker, non-specific staining was removed with destaining buffer (250 g chloral hydrate in 100 ml H_2_O). Finally, plant tissues were kept in 50% (v/v) glycerol for imaging. For the measurement of electrolyte leakage, ten first or second leaves from independent plants were harvested at the indicated time points and transferred to a 15 ml tube containing 6 ml water purified using a Milli-Q integral 5 water purification system (Millipore). After a 6 h-incubation at room temperature on a horizontal shaker, conductivity of the solution was measured with an Orion Star A212 conductivity meter (Thermo Scientific). This experiment was repeated three times with similar results.

### Determination of maximum photochemical efficiency of Photosystem II (Fv/Fm)

The Fv/Fm was determined with a FluorCam system (FC800-C/1010GFP; Photon Systems Instruments) containing a CCD camera and an irradiation system according to the instrument manufacturer's instructions.

### Induced resistance

Induced resistance assays were performed as described previously^70^ with slight modifications. Briefly, 5 μM solutions of the PNP-A peptide or its scrambled peptide control were infiltrated with a needleless syringe into the abaxial side of leaves of five-week-old short-day-grown Arabidopsis plants 6 h prior to bacterial infection (*Pseudomonas syringae* pv. *tomato* DC3000, 10^5^ cfu/ml). Bacterial growth was determined three days after inoculation by plating 1:10 serial dilutions of leaf extracts; plates were incubated at 28 °C for 2 days before bacterial cfu were counted. For each treatment, 7 mm leaf discs of three leaves from four independent plants were used.

## Acknowledgements

We thank the Core Facility of Proteomics, Shanghai Center for Plant Stress Biology (PSC) for carrying out mass spectrometry. We thank Junghee Lee for critical comments on the manuscript. This research was supported by the 100-Talents Program from the Chinese Academy of Sciences (CAS) to C.K., by National Natural Science Foundation of China (NSFC) Grant 31570264 to C.K., and by the Strategic Priority Research Program XDB27040102 from the CAS to C.K..

## Author contributions

K.P.L., K.L., E.Y.K., J.D., L.M.P., Y.L., H.D., R.L., and Z.L. conducted the experiments; K.P.L., K.L., E.Y.K., J.D., L.M.P., R.L.D., and C.K. designed the research; K.P.L., K.L., E.Y.K., J.D., L.M.P., R.L.D., and C.K. analyzed the data; K.P.L. and C.K. wrote the manuscript. All authors reviewed and edited the manuscript.

## Competing financial interests

The authors declare no competing financial interests.

## Additional information

**Supplementary Figure 1. *PNP-A* is highly upregulated in *lsd1*.** (**a**) The transcript levels of *PNP-A* in 17-d and 19-d-old plants of wild-type (WT) and *lsd1* grown under continuous light condition (CL) were obtained from our previous RNA-Seq analysis^27^. cpm: count per million. (**b**) The transcript levels of *PNP-A* shown in **a** were confirmed by qRT-PCR. (**c**) WT and *npr1* plants grown under CL were sprayed with a 0.5 mM solution of SA (+ SA) or with distilled water (− SA), and leaf samples were harvested at 12 hrs after the treatment. Expression level of *PNP-A* was examined using qRT-PCR. (**d**) Expression levels of *PNP-A* in WT, *lsd1* and *lsd1 npr1* (*l*/*n1*) grown under CL were analyzed by qRT-PCR at the indicated time points. For the qRT-PCR analyses in **b**, **c**, and **d**, *ACT2* was used as an internal standard. The data represent the means of three independent biological replicates. Error bars indicate standard deviation. Lowercase letters indicate statistically significant differences between mean values (*P* < 0.01, one-way ANOVA with post-hoc Tukey’s HSD test).

**Supplementary Figure 2. Two independent wild-type transgenic lines overexpressing PNP-A.** Transcript levels of *PNP-A* in two independent *lsd1* plants overexpressing *GFP*-tagged *PNP-A* (*PNP-A-GFP*) under the control of the *CaMV 35S* promoter. Semiquantitative RT-PCR (**a**) and quantitative RT-PCR (**b**) were carried out with total RNAs isolated from the 16-d-old CL-grown plants. *ACT2* was used as an internal standard. The data in **b** represent the means of three independent biological replicates. Error bars indicate standard deviation. Lowercase letters indicate statistically significant differences between mean values (*P* < 0.01, one-way ANOVA with post-hoc Tukey’s HSD test).

**Supplementary Figure 3. Effect of loss of or overexpression of PNP-A on*lsd1* RCD.** WT, *lsd1*, *lsd1 pnp-A* and two PNP-A overexpression lines were grown under CL and the emergence and spread of RCD were monitored at the indicated time points. The images of representative plants are shown at the same scale.

**Supplementary Figure 4. PNP-A represses SA biosynthesis genes and SA-responsive genes.** WT, *lsd1*, *lsd1 pnp-A* and two PNP-A overexpression lines were grown under CL and expression levels of genes involved in SA biosynthesis (*ICS1*, *EDS1*, and *PAD4*) and SA response (*PR1* and *PR2*) were examined by qRT-PCR at the indicated time points. *ACT2* was used as an internal standard. The data represent the means of three independent biological replicates. Error bars indicate standard deviation.

**Supplementary Figure 5. Schematic illustration of domain structure of *Arabidopsis* PNP-A (AtPNP-A).** Signal peptide (1 to 29 amino acids) is responsible for targeting to extracellular space. Amino acids 36 to 69 indicate the active region of PNP-A that has significant biological activity. The amino acid sequences of N-terminally biotinylated active region of synthetic AtPNP-A (Active) and scrambled peptides are represented. Red capital letter “C” indicates two cystein residues that form a disulfide bond, as indicated in a previous study^15^.

**Supplementary Table 1. List of proteins interacting PNP-A synthetic peptides.** Pull-down with N-terminal biotinylated PNP-A synthetic peptide coupled to mass spectrometry analysis (PNP-A pull-down/MS) was performed with total proteins extracted from *lsd1* mutant plants grown under continuous light on MS medium for 20 days. The pull-down/MS with N-terminally biotinylated scrambled PNP-A or without the synthetic peptides were also performed for negative controls. The PNP-A pull-down/MS repeated two times with independent biological samples. A total of 66 proteins were present in both biological replicates from the PNP-A pull-down/MS, but were absent in the negative controls.

**Supplementary Table 2. List of primer sets used in this study.**

## References

1. Van Norman J. M., Breakfield N. W. & Benfey P. N. Intercellular communication during plant development. Plant Cell. 23, 855–864(2011).

2. Lease K. A. & Walker J. C. The Arabidopsis unannotated secreted peptide database, a resource for plant peptidomics. Plant Physiol. 142, 831–838 (2006).

3. Matsubayashi Y. Posttranslationally Modified Small-Peptide Signals in Plants. Annual Review of Plant Biology, Vol 65, 385–413 (2014).

4. Butenko M. A., Vie A. K., Brembu T., Aalen R. B. & Bones A. M. Plant peptides in signalling: looking for new partners. Trends Plant Sci. 14, 255–263 (2009).

5. Matsubayashi Y. Post-Translational Modifications in Secreted Peptide Hormones in Plants. Plant Cell Physiol. 52, 5–13 (2011).

6. Murphy E., Smith S. & De Smet I. Small Signaling Peptides in Arabidopsis Development: How Cells Communicate Over a Short Distance. Plant Cell. 24, 3198–3217 (2012).

7. Motomitsu A., Sawa S. & Ishida T. Plant peptide hormone signalling. Essays Biochem. 58, 115–131 (2015).

8. Schardon K. et al. Precursor processing for plant peptide hormone maturation by subtilisin-like serine proteinases. Science. 354, 1594–1597 (2016).

9. Shiu S. H. & Bleecker A. B. Receptor-like kinases from Arabidopsis form a monophyletic gene family related to animal receptor kinases. P Natl Acad Sci USA. 98, 10763–10768 (2001).

10. Wang G. D. et al. A genome-wide functional investigation into the roles of receptor-like proteins in Arabidopsis. Plant Physiol. 147, 503–517 (2008).

11. Hirakawa Y., Torii K. U. & Uchida N. Mechanisms and Strategies Shaping Plant Peptide Hormones. Plant Cell Physiol. 58, 1313–1318 (2017).

12. Ludidi N. N., Heazlewood J. L., Seoighe C., Irving H. R. & Gehring C. A. Expansin-like molecules: Novel functions derived from common domains. J Mol Evol. 54, 587–594 (2002).

13. Pharmawati M., Maryani M. M., Nikolakopoulos T., Gehring C. A. & Irving H. R. Cyclic GMP modulates stomatal opening induced by natriuretic peptides and immunoreactive analogues. Plant Physiol Bioch. 39, 385–394 (2001).

14. Wang Y. H., Gehring C., Cahill D. M. & Irving H. R. Plant natriuretic peptide active site determination and effects on cGMP and cell volume regulation. Funct Plant Biol. 34, 645–653 (2007).

15. Turek I. & Gehring C. The plant natriuretic peptide receptor is a guanylyl cyclase and enables cGMP-dependent signaling. Plant Mol Biol. 91, 275–286 (2016).

16. Gehring C. & Turek I. S. Cyclic Nucleotide Monophosphates and Their Cyclases in Plant Signaling. Front Plant Sci. 8, (2017).

17. Potter L. R. & Hunter T. Guanylyl cyclase-linked natriuretic peptide receptors: Structure and regulation. J Biol Chem. 276, 6057–6060 (2001).

18. Morse M., Pironcheva G. & Gehring C. AtPNP-A is a systemically mobile natriuretic peptide immunoanalogue with a role in Arabidopsis thaliana cell volume regulation. Febs Lett. 556, 99–103 (2004).

19. Gottig N. et al. Xanthomonas axonopodis pv. citri uses a plant natriuretic peptide-like protein to modify host homeostasis. P Natl Acad Sci USA. 105, 18631–18636 (2008).

20. Ficarra F. A., Grandellis C., Garavaglia B. S., Gottig N. & Ottado J. Bacterial and plant natriuretic peptides improve plant defence responses against pathogens. Mol Plant Pathol. 19, 801–811 (2018).

21. Ruzvidzo O., Donaldson L., Valentine A. & Gehring C. The Arabidopsis thaliana natriuretic peptide AtPNP-A is a systemic regulator of leaf dark respiration and signals via the phloem. J Plant Physiol. 168, 1710–1714 (2011).

22. Ludidi N. et al. A recombinant plant natriuretic peptide causes rapid and spatially differentiated K+, Na+ and H+ flux changes in Arabidopsis thaliana roots. Plant Cell Physiol. 45, 1093–1098 (2004).

23. Wang Y. H., Donaldson L., Gehring C. & Irving H. R. Plant natriuretic peptides: control of synthesis and systemic effects. Plant Signal Behav. 6, 1606–1608 (2011).

24. Wang Y. H., Gehring C. & Irving H. R. Plant Natriuretic Peptides are Apoplastic and Paracrine Stress Response Molecules. Plant Cell Physiol. 52, 837–850 (2011).

25. Cao H., Glazebrook J., Clarke J. D., Volko S. & Dong X. N. The Arabidopsis NPR1 gene that controls systemic acquired resistance encodes a novel protein containing ankyrin repeats. Cell. 88, 57–63 (1997).

26. Zhang Y. L., Fan W. H., Kinkema M., Li X. & Dong X. N. Interaction of NPR1 with basic leucine zipper protein transcription factors that bind sequences required for salicylic acid induction of the PR-1 gene. P Natl Acad Sci USA. 96, 6523–6528 (1999).

27. Lv R. et al. Uncoupled Expression of Nuclear and Plastid Photosynthesis-associated Genes Contributes to Cell Death in a Lesion Mimic Mutant. Plant Cell. (2019).

28. Dietrich R. A. et al. Arabidopsis Mutants Simulating Disease Resistance Response. Cell. 77, 565–577 (1994).

29. Jabs T., Dietrich R. A. & Dangl J. L. Initiation of runaway cell death in an Arabidopsis mutant by extracellular superoxide. Science. 273, 1853–1856 (1996).

30. Dietrich R. A., Richberg M. H., Schmidt R., Dean C. & Dangl J. L. A novel zinc finger protein is encoded by the arabidopsis LSD1 gene and functions as a negative regulator of plant cell death. Cell. 88, 685–694 (1997).

31. Kliebenstein D. J., Dietrich R. A., Martin A. C., Last R. L. & Dangl J. L. LSD1 regulates salicylic acid induction of copper zinc superoxide dismutase in Arabidopsis thaliana. Mol Plant Microbe In. 12, 1022–1026 (1999).

32. Rusterucci C., Aviv D. H., Holt B. F., Dangl J. L. & Parker J. E. The disease resistance signaling components EDS1 and PAD4 are essential regulators of the cell death pathway controlled by LSD1 in arabidopsis. Plant Cell. 13, 2211–2224 (2001).

33. Aviv D. H. et al. Runaway cell death, but not basal disease resistance, in Isd1 is SA- and NIM1/NPR1-dependent. Plant J. 29, 381–391 (2002).

34. Wang D., Amornsiripanitch N. & Dong X. N. A genomic approach to identify regulatory nodes in the transcriptional network of systemic acquired resistance in plants. Plos Pathog. 2, 1042–1050 (2006).

35. Schwartz D. et al. Ser-Leu-Arg-Arg-atriopeptin III: the major circulating form of atrial peptide. Science. 229, 397–400 (1985).

36. Koller K. J. & Goeddel D. V. Molecular-Biology of the Natriuretic Peptides and Their Receptors. Circulation. 86, 1081–1088 (1992).

37. Vicente M. R. S. & Plasencia J. Salicylic acid beyond defence: its role in plant growth and development. J Exp Bot. 62, 3321–3338 (2011).

38. Karasov T. L., Chae E., Herman J. J. & Bergelson J. Mechanisms to Mitigate the Trade-Off between Growth and Defense. Plant Cell. 29, 666–680 (2017).

39. Scott J. D. Cyclic Nucleotide-Dependent Protein-Kinases. Pharmacol Therapeut. 50, 123–145 (1991).

40. Durner J., Wendehenne D. & Klessig D. F. Defense gene induction in tobacco by nitric oxide, cyclic GMP, and cyclic ADP-ribose. P Natl Acad Sci USA. 95, 10328–10333 (1998).

41. Donaldson L., Ludidi N., Knight M. R., Gehring C. & Denby K. Salt and osmotic stress cause rapid increases in Arabidopsis thaliana cGMP levels. Febs Lett. 569, 317–320 (2004).

42. Hoshi T. Regulation of Voltage-Dependence of the Kat1 Channel by Intracellular Factors. J Gen Physiol. 105, 309–328 (1995).

43. Leng Q., Mercier R. W., Yao W. Z. & Berkowitz G. A. Cloning and first functional characterization of a plant cyclic nucleotide-gated cation channel. Plant Physiol. 121, 753–761 (1999).

44. Neill S. J., Desikan R. & Hancock J. T. Nitric oxide signalling in plants. New Phytol. 159, 11–35 (2003).

45. Neill S. et al. Nitric oxide, stomatal closure, and abiotic stress. J Exp Bot. 59, 165–176 (2008).

46. Penson S. P. et al. cGMP is required for gibberellic acid-induced gene expression in barley aleurone. Plant Cell. 8, 2325–2333 (1996).

47. Zhao Y. C., Qi Z. & Berkowitz G. A. Teaching an Old Hormone New Tricks: Cytosolic Ca2+ Elevation Involvement in Plant Brassinosteroid Signal Transduction Cascades. Plant Physiol. 163, 555–565 (2013).

48. Bowler C., Yamagata H., Neuhaus G. & Chua N. H. Phytochrome Signal-Transduction Pathways Are Regulated by Reciprocal Control Mechanisms. Gene Dev. 8, 2188–2202 (1994).

49. Krogh A., Larsson B., von Heijne G. & Sonnhammer E. L. L. Predicting transmembrane protein topology with a hidden Markov model: Application to complete genomes. J Mol Biol. 305, 567–580 (2001).

50. Kall L., Krogh A. & Sonnhammer E. L. L. A combined transmembrane topology and signal peptide prediction method. J Mol Biol. 338, 1027–1036 (2004).

51. Marchler-Bauer A. et al. CDD/SPARCLE: functional classification of proteins via subfamily domain architectures. Nucleic Acids Res. 45, D200–D203 (2017).

52. Finn R. D. et al. InterPro in 2017-beyond protein family and domain annotations. Nucleic Acids Res. 45, D190–D199 (2017).

53. Ludidi N. & Gehring C. Identification of a novel protein with guanylyl cyclase activity in Arabidopsis thaliana. J Biol Chem. 278, 6490–6494 (2003).

54. Dempsey D. A., Shah J. & Klessig D. F. Salicylic acid and disease resistance in plants. Crit Rev Plant Sci. 18, 547–575 (1999).

55. Meier S. et al. Co-expression and promoter content analyses assign a role in biotic and abiotic stress responses to plant natriuretic peptides. Bmc Plant Biol. 8, (2008).

56. Mateo A. et al. LESION SIMULATING DISEASE 1 is required for acclimation to conditions that promote excess excitation energy. Plant Physiol. 136, 2818–2830 (2004).

57. Muhlenbock P. et al. Chloroplast signaling and LESION SIMULATING DISEASE1 regulate crosstalk between light acclimation and immunity in Arabidopsis. Plant Cell. 20, 2339–2356 (2008).

58. Huang X., Li Y., Zhang X., Zuo J. & Yang S. The Arabidopsis LSD1 gene plays an important role in the regulation of low temperature-dependent cell death. New Phytol. 187, 301–312 (2010).

59. Chai T., Zhou J., Liu J. & Xing D. LSD1 and HY5 antagonistically regulate red light induced-programmed cell death in Arabidopsis. Front Plant Sci. 6, 292 (2015).

60. Wituszynska W. et al. Lesion simulating disease 1 and enhanced disease susceptibility 1 differentially regulate UV-C-induced photooxidative stress signalling and programmed cell death in Arabidopsis thaliana. Plant Cell Environ. 38, 315–330 (2015).

61. Turek I., Marondedze C., Wheeler J. I., Gehring C. & Irving H. R. Plant natriuretic peptides induce proteins diagnostic for an adaptive response to stress. Front Plant Sci. 5, (2014).

62. Torres M. A., Jones J. D. & Dangl J. L. Pathogen-induced, NADPH oxidase-derived reactive oxygen intermediates suppress spread of cell death in Arabidopsis thaliana. Nat Genet. 37, 1130–1134 (2005).

63. Jeong S., Trotochaud A. E. & Clark S. E. The Arabidopsis CLAVATA2 gene encodes a receptor-like protein required for the stability of the CLAVATA1 receptor-like kinase. Plant Cell. 11, 1925–1933 (1999).

64. Nadeau J. A. & Sack F. D. Control of stomatal distribution on the Arabidopsis leaf surface. Science. 296, 1697–1700 (2002).

65. Nakagawa T. et al. Improved gateway binary vectors: High-performance vectors for creation of fusion constructs in Transgenic analysis of plants. Biosci Biotech Bioch. 71, 2095–2100 (2007).

66. Clough S. J. & Bent A. F. Floral dip: a simplified method for Agrobacterium-mediated transformation of Arabidopsis thaliana. Plant J. 16, 735–743 (1998).

67. Livak K. J. & Schmittgen T. D. Analysis of relative gene expression data using real-time quantitative PCR and the 2(T)(-Delta Delta C) method. Methods. 25, 402–408 (2001).

68. Lu Q. et al. Arabidopsis homolog of the yeast TREX-2 mRNA export complex: components and anchoring nucleoporin. Plant J. 61, 259–270 (2010).

69. Wang L. S. et al. Singlet oxygen- and EXECUTER1-mediated signaling is initiated in grana margins and depends on the protease FtsH2. P Natl Acad Sci USA. 113, E3792–E3800 (2016).

70. Lozano-Duran R. et al. The transcriptional regulator BZR1 mediates trade-off between plant innate immunity and growth. Elife. 2, e00983 (2013).

