## Supplementary Figures for "PLANT NATRIURETIC PEPTIDE A antagonizes salicylic acid-primed cell death"

### Supplementary Fig. 1

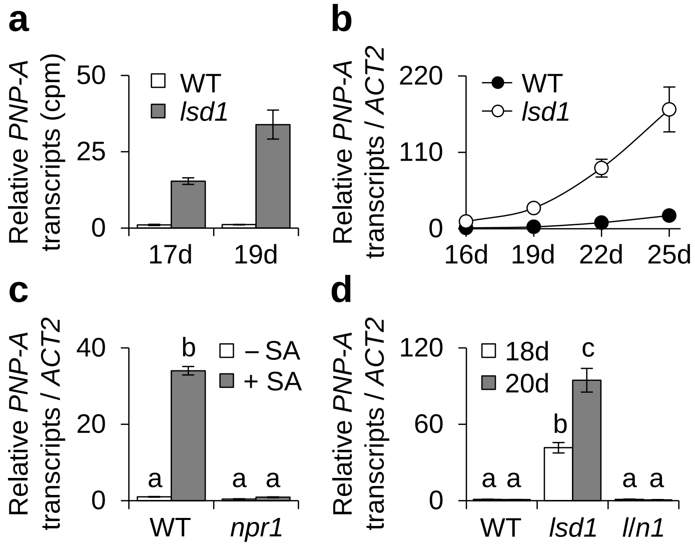


**Supplementary Figure 1. *PNP-A* is highly upregulated in *lsd1*.** (**a**) The transcript levels of *PNP-A* in 17-d and 19-d-old plants of wild-type (WT) and *lsd1* grown under continuous light condition (CL) were obtained from our previous RNA-Seq analysis[29]. cpm: count per million. (**b**) The transcript levels of *PNP-A* shown in **a** were confirmed by qRT-PCR. (**c**) WT and *npr1* plants grown under CL were sprayed with a 0.5 mM solution of SA (+ SA) or with distilled water (- SA), and leaf samples were harvested at 12 hrs after the treatment. Expression level of *PNP-A* was examined using qRT-PCR. (**d**) Expression levels of *PNP-A* in WT, *lsd1* and *lsd1 npr1* (*l/n1*) grown under CL were analyzed by qRT-PCR at the indicated time points. For the qRT-PCR analyses in **b**, **c**, and **d**, *ACT2* was used as an internal standard. The data represent the means of three independent biological replicates. Error bars indicate standard deviation. Lowercase letters indicate statistically significant differences between mean values ($P < 0.01$, one-way ANOVA with post-hoc Tukey's HSD test).

### Supplementary Fig. 2

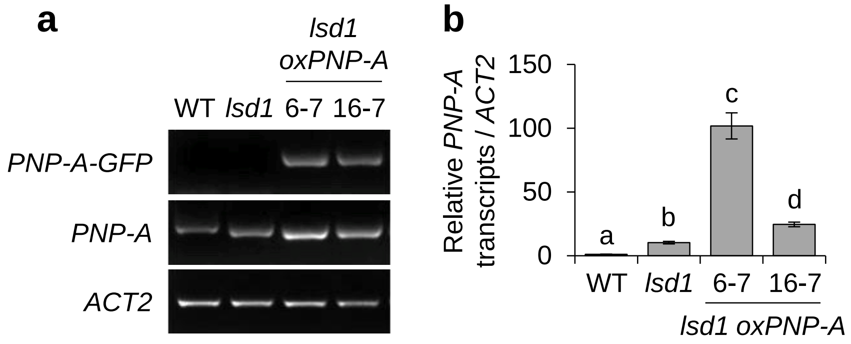


**Supplementary Figure 2. Two independent wild-type transgenic lines overexpressing PNP-A.** Transcript levels of *PNP-A* in two independent *lsd1* plants overexpressing *GFP*-tagged *PNP-A* (*PNP-A-GFP*) under the control of the *CaMV 35S* promoter. Semiquantitative RT-PCR (**a**) and quantitative RT-PCR (**b**) were carried out with total RNAs isolated from the 16-d-old CL-grown plants. *ACT2* was used as an internal standard. The data in **b** represent the means of three independent biological replicates. Error bars indicate standard deviation. Lowercase letters indicate statistically significant differences between mean values ($P < 0.01$, one-way ANOVA with post-hoc Tukey's HSD test).

#### Supplementary Fig. 3

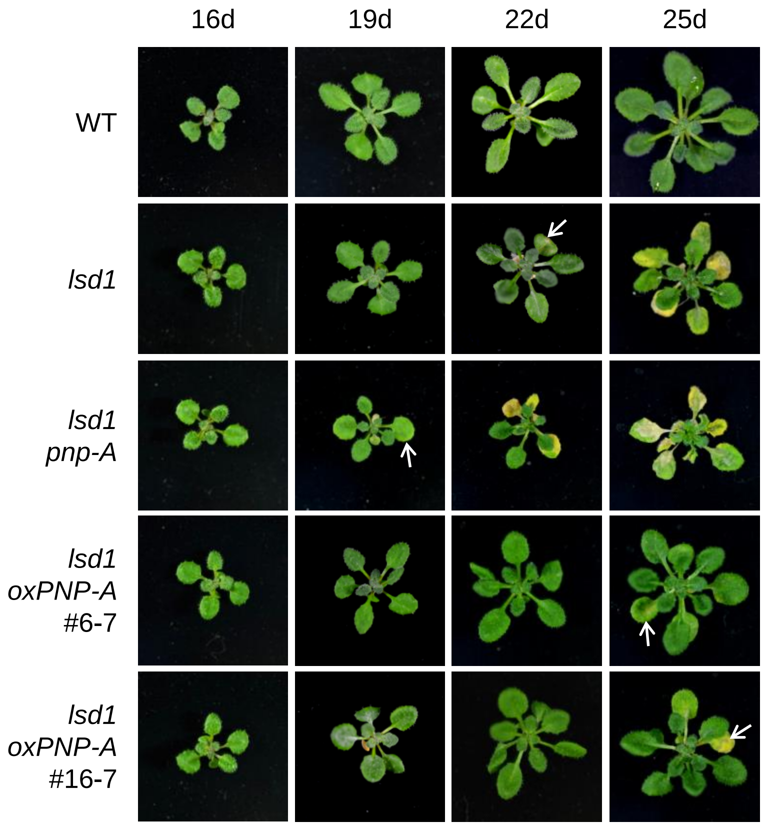


**Supplementary Figure 3**. **Effect of loss of or overexpression of PNP-A on *lsd1* RCD.** WT, *lsd1*, *lsd1 pnp-A* and two PNP-A overexpression lines were grown under CL and the emergence and spread of RCD were monitored at the indicated time points. The images of representative plants are shown at the same scale.

### Supplementary Fig. 4

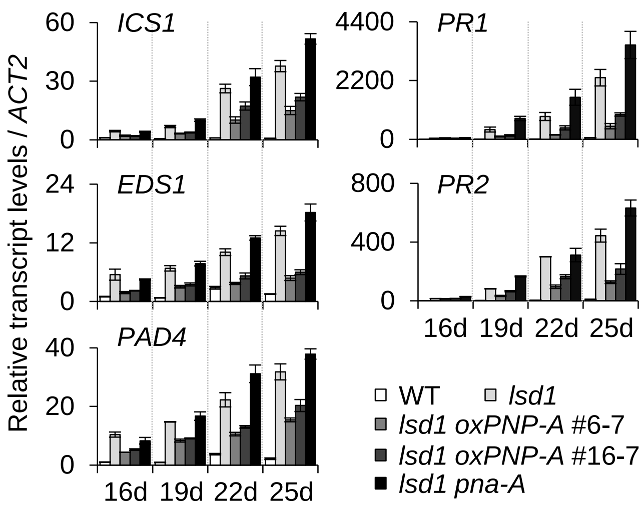


**Supplementary Figure 4. PNP-A represses SA biosynthesis genes and SA-responsive genes.** WT, *lsd1*, *lsd1 pnp-A* and two PNP-A overexpression lines were grown under CL and expression levels of genes involved in SA biosynthesis (*ICS1*, *EDS1*, and *PAD4*) and SA response (*PR1* and *PR2*) were examined by qRT-PCR at the indicated time points. *ACT2* was used as an internal standard. The data represent the means of three independent biological replicates. Error bars indicate standard deviation.

### Supplementary Fig. 5

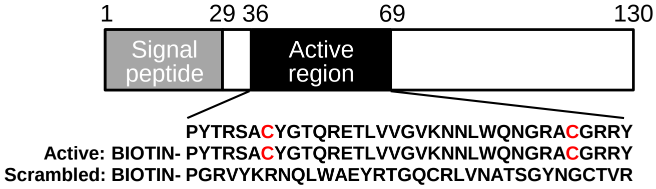


**Supplementary Figure 5. Schematic illustration of domain structure of Arabidopsis PNP-A (AtPNP-A).** Signal peptide (1 to 29 amino acids) is responsible for targeting to extracellular space. Amino acids 36 to 69 indicate the active region of PNP-A that has significant biological activity. The amino acid sequences of N-terminally biotinylated active region of synthetic AtPNP-A (Active) and scrambled peptides are represented. Red capital letter “C” indicates two cystein residues that form a disulfide bond, as indicated in a previous study[15].
