## Supplementary Tables for "PLANT NATRIURETIC PEPTIDE A antagonizes salicylic acid-primed cell death"

**Supplementary Table 1. List of proteins interacting PNP-A synthetic peptides.** Pull-down with N-terminal biotinylated PNP-A synthetic peptide coupled to mass spectrometry analysis (PNP-A pull-down/MS) was performed with total proteins extracted from *lsd1* mutant plants grown under continuous light on MS medium for 20 days. The pull-down/MS with N-terminally biotinylated scrambled PNP-A or without the synthetic peptides were also performed for negative controls. The PNP-A pull-down/MS repeated two times with independent biological samples. A total of 66 proteins were present in both biological replicates from the PNP-A pull-down/MS, but were absent in the negative controls.

| **Nr.** | **Gene ID** | **Protein name & Description** | **Nr. of peptide** | |
| --- | --- | --- | --- | --- |
|  |  |  | **1st** | **2nd** |
| 1 | AT1G07320 | RPL4, ribosomal protein L4 | 2 | 5 |
| 2 | AT1G10670 | ACLA-1, ATP-citrate lyase A-1 | 1 | 1 |
| 3 | AT1G11840 | GLX1, Ni+ dependent glyoxalase I homolog | 4 | 2 |
| 4 | AT1G13190 | RNA-binding (RRM/RBD/RNP motifs) family protein | 5 | 3 |
| 5 | AT1G23290 | RPL27A, ribosomal protein L18e/L15 superfamily protein | 3 | 4 |
| 6 | AT1G23860 | RSZP21, SRZ-21, SRZ21, RS-containing zinc finger protein 21 | 4 | 1 |
| 7 | AT1G26110 | DCP5, decapping 5; required for mRNA decapping, | 4 | 3 |
| 8 | AT1G26230 | TCP-1/cpn60 chaperonin family protein | 1 | 1 |
| 9 | AT1G36240 | Ribosomal protein L7Ae/L30e/S12e/Gadd45 family protein | 4 | 1 |
| 10 | AT1G43700 | SUE3, VIP1, VIRE2-interacting protein 1 | 3 | 1 |
| 11 | AT1G48030 | mtLPD1, mitochondrial lipoamide dehydrogenase 1 | 1 | 1 |
| 12 | AT1G55170 | Unknown protein | 6 | 4 |
| 13 | AT1G55810 | UKL3, uridine kinase-like 3 | 10 | 15 |
| 14 | AT1G56190 | Phosphoglycerate kinase family protein | 1 | 1 |
| 15 | AT1G64390 | GH9C2, glycosyl hydrolase 9C2 | 2 | 2 |
| 16 | AT1G67170 | Unknown protein | 3 | 2 |
| 17 | AT1G67750 | Pectate lyase family protein | 1 | 1 |
| 18 | AT1G70180 | Sterile alpha motif (SAM) domain-containing protein | 3 | 2 |
| 19 | AT1G70710 | GH9B1, glycosyl hydrolase 9B1 | 1 | 3 |
| 20 | AT1G76300 | SmD3, snRNP core protein SMD3 | 1 | 1 |
| 21 | AT1G77940 | Ribosomal protein L7Ae/L30e/S12e/Gadd45 family protein | 1 | 2 |
| 22 | AT1G78630 | EMB1473, Ribosomal protein L13 family protein | 3 | 3 |
| 23 | AT1G79550 | PGK, cytosolic phosphoglycerate kinase | 1 | 2 |
| **24** | **AT2G18660** | **PNP-A, plant natriuretic peptide A** | **6** | **6** |
| 25 | AT2G18740 | Small nuclear ribonucleoprotein family protein | 3 | 3 |
| 26 | AT2G20450 | Ribosomal protein L14 | 1 | 1 |
| 27 | AT2G27530 | PGY1, Ribosomal protein L1p/L10e family | 4 | 1 |
| 28 | AT2G29140 | APUM3, PUM3, pumilio 3 | 1 | 1 |
| 29 | AT2G33410 | RNA-binding (RRM/RBD/RNP motifs) family protein | 3 | 2 |
| 30 | AT2G34160 | Alba DNA/RNA-binding protein | 1 | 3 |
| 31 | AT2G40650 | PRP38 family protein | 4 | 2 |
| 32 | AT3G06720 | IMPA1, importin alpha isoform 1 | 3 | 5 |
| 33 | AT3G13224 | RNA-binding (RRM/RBD/RNP motifs) family protein | 5 | 3 |
| 34 | AT3G13580 | Ribosomal protein L30/L7 family protein | 1 | 1 |
| 35 | AT3G14750 | Unknown protein | 3 | 2 |
| **Nr.** | **Gene ID** | **Protein name & Description** | **Nr. of peptide** | |
|  |  |  | **1st** | **2nd** |
| 36 | AT3G18035 | HON4, A linker histone like protein | 7 | 7 |
| 37 | AT3G18390 | EMB1865, CRS1 / YhbY (CRM) domain-containing protein | 2 | 4 |
| 38 | AT3G18640 | Zinc finger C-x8-C-x5-C-x3-H type family protein | 4 | 2 |
| 39 | AT3G19130 | RBP47B, RNA-binding protein 47B | 4 | 1 |
| 40 | AT3G43670 | Copper amine oxidase family protein | 1 | 6 |
| 41 | AT3G45830 | Unknown protein | 2 | 6 |
| 42 | AT3G49120 | PRX34, PRXCB, peroxidase CB | 2 | 1 |
| 43 | AT3G49430 | SRp34a, serine/arginine-rich protein splicing factor 34A | 2 | 7 |
| 44 | AT3G54230 | SUA, suppressor of abi3-5 | 13 | 6 |
| 45 | AT3G62870 | Ribosomal protein L7Ae/L30e/S12e/Gadd45 family protein | 6 | 2 |
| 46 | AT4G02840 | Small nuclear ribonucleoprotein family protein | 1 | 1 |
| 47 | AT4G10110 | RNA-binding (RRM/RBD/RNP motifs) family protein | 1 | 2 |
| 48 | AT4G14880 | OLD3, Onset of leaf death 3 | 4 | 2 |
| 49 | AT4G17060 | FIP2, FRIGIDA interacting protein 2 | 4 | 4 |
| 50 | AT4G23600 | JR2, JA responsive 2 | 2 | 2 |
| 51 | AT4G24770 | RBP31, 31-kDa RNA binding protein | 4 | 3 |
| 52 | AT4G26110 | NAP1;1, nucleosome assembly protein1;1 | 2 | 1 |
| 53 | AT4G28990 | RNA-binding protein-related | 3 | 5 |
| 54 | AT4G31180 | IBI1, Impaired in BABA-induced disease immunity 1 | 2 | 2 |
| 55 | AT5G02450 | Ribosomal protein L36e family protein | 3 | 1 |
| 56 | AT5G02490 | HSP70, Heat shock protein 70 | 2 | 1 |
| **57** | **AT5G12940** | **Leucine-rich repeat (LRR) family protein** | **2** | **7** |
| 58 | AT5G13590 | Unknown protein | 2 | 8 |
| 59 | AT5G19550 | ASP2, Aspartate aminotransferase 2 | 1 | 2 |
| 60 | AT5G19960 | RNA-binding (RRM/RBD/RNP motifs) family protein | 3 | 4 |
| 61 | AT5G38600 | Proline-rich spliceosome-associated (PSP) family | 6 | 3 |
| 62 | AT5G40490 | RNA-binding (RRM/RBD/RNP motifs) family protein | 8 | 6 |
| 63 | AT5G56500 | TCP-1/cpn60 chaperonin family protein | 1 | 1 |
| 64 | AT5G58470 | TAF15b, TBP-associated factor 15B | 2 | 3 |
| 65 | AT5G65220 | Ribosomal L29 family protein | 1 | 3 |
| 66 | ATCG00660 | RPL20, ribosomal protein L20 | 2 | 3 |

**Supplementary Table 2.** **List of primer sets used in this study.**

| **Gene ID** | **Gene**  **name** | **Mutant**  **alleles** | **Primer sequence (5' to 3')** | **Primer**  **length** | **Size**  **(bp)** | **Used**  **for** |
| --- | --- | --- | --- | --- | --- | --- |
| At3g18780 | *ACT2* | *-* | F: TATGTATGTCGCCATCCAA | 19 | 76 | qRT-PCR |
|  |  |  | R: ACCAGAATCCAGCACAATA | 19 |  |  |
| At2g18660 | *PNP-A* | - | F: TCTATTACGACCCTCCCTACAC | 22 | 80 |  |
|  |  |  | R: ATTGTTCTTGACTCCGACTACC | 22 |  |  |
| At1g74710 | *ICS1* | *-* | F: GTACAGTGGAGACAAGGACTATG | 23 | 102 |  |
|  |  |  | R: ACGACGGAGGTTGAGATTTG | 20 |  |  |
| At2g14610 | *PR1* | *-* | F: AGTGAGGTGTAACAATGGT | 19 | 82 |  |
|  |  |  | R: TTAGTATGGCTTCTCGTTCA | 20 |  |  |
| At3g57260 | *PR2* | *-* | F: CCACCAATGTTGATGATTCT | 20 | 83 |  |
|  |  |  | R: CCGTAGCATACTCCGATT | 18 |  |  |
| At3g48090 | *EDS1* | *-* | F: CCATACGAGGAAGTTGAGGTAAG | 23 | 110 |  |
|  |  |  | R: AACGTTGAACCCTCCAGAAATA | 22 |  |  |
| At3g52430 | *PAD4* | *-* | F: GCACAACGCAAGATACTT | 18 | 63 |  |
|  |  |  | R: ATGTACGGCCCTGTGTCTTC | 18 |  |  |
| At2g18660 | *PNP-A* | - | F: GCGTACCGTAGACGTGAAGG | 20 | 225 | RT-PCR |
|  |  |  | R: TTTCCGTATGTTGCATCACC | 20 |  |  |
| At3g18780 | *ACT2* | - | F: GGCTCCTCTTAACCCAAAGG | 20 | 265 |  |
|  |  |  | R: CAGTAAGGTCACGTCCAGCA | 20 |  |  |
|  | *PNP-A-GFP* | - | F: GCAATAAGCCACATAATGCAG | 21 | 255 |  |
|  |  |  | R: CTGACCAATTAGGAGTCAGAAACTAC | 26 |  |  |
| At2g18660 | *PNP-A* | SALK_000951 | LP: TTTCCGGATATCCGAAATTTC | 21 | 1016 | Genotyping |
|  |  |  | RP: ATGTGATTTAGGGTCTGCGTG | 21 |  |  |
| At3g56710 | *PNPAR* | GABIseq255A11.1  (*pnpar-1*) | LP: AGCCACTTGAATGTTGTCCC | 20 | 1110 |  |
|  |  |  | RP: TATTCCTGAGATCCAAATGCG | 21 |  |  |
| At2g41180 | *PNPAR* | SALKseq6749.3  (*pnpar-2*) | LP: CTAGACTGTTGCAAAGGCTGG | 21 | 1140 |  |
|  |  |  | RP: TTTCATTTGGTTCGAGGTGTC | 21 |  |  |
| At1g33612 | *PNP-R1* | GABI_180G04 | LP: ATTACTTGACTGGGTTTGCCC | 25 | 1191 |  |
|  |  |  | RP: CGAAGAATAAGTACTCGGGGG | 25 |  |  |
